# Improved coherence measure for EEG-based speech tracking in hearing impaired listeners

**DOI:** 10.1101/2023.12.05.570227

**Authors:** Oskar Keding, Emina Alickovic, Martin Skoglund, Maria Sandsten

## Abstract

In the literature, auditory attention is explored through neural speech tracking, primarily entailing modeling and analyzing electroencephalography (EEG) responses to natural speech via linear filtering. Our study takes a novel approach, introducing an enhanced coherence estimation technique that employs multitapers to assess the strength of neural speech tracking. This enables effective discrimination between attended and ignored speech. To mitigate the impact of colored noise in EEG, we address two biases – overall coherence-level bias and spectral peak-shifting bias. In a listening study involving 32 participants with hearing impairment, tasked with attending to competing talkers in background noise, our coherence-based method effectively discerns EEG representations of attended and ignored speech. We comprehensively analyze frequency bands, individual frequencies, and EEG channels. Frequency bands of importance are shown to be delta, theta and alpha, as well as the central EEG channels. Lastly, we showcase coherence differences across different noise reduction settings implemented in hearing aids, underscoring our method’s potential to objectively assess auditory attention and enhance hearing aid efficacy.

## 1 Introduction

Listening is a biological process that engages the entire auditory system. The process, from sound pressures to neural activations, includes (non-linear) transforms of the peripheral auditory system as well as complex processing within the central auditory system. The listening process also affects the electrical activity of the brain, which can be measured on scalp-level using electroencephalography (EEG). Linear filter estimation, referred to as temporal response functions (TRFs) [1–3], has been shown to capture the relations between the auditory system and EEG signatures [4]. State-of-the art methods employ TRFs to analyze phonemes [5–7] and semantic content [8, 9]. The TRF methods are also particularly valuable for decoding auditory attention, specifically in identifying the attended talker from EEG signals in challenging listening situations like cocktail-party environments [10]. Effectively decoding auditory attention to distinguish and enhance attended speech in multi-talker situations holds significance for hearing aid (HA) applications [11–14].

The mentioned studies, along with others, provide evidence that neural speech processing exhibits a linear component. Specifically, there are robust linear relationships between the speech envelope and the concurrent EEG signatures. Hence, spectral coherence analysis between the speech envelope and EEG signals can confidently identify and analyze the underlying system to a significant degree, as demonstrated in [15, 16]. Spectral coherence is a measure of the linear coupling between two signals, calculated as the cross-spectrum normalized by the respective auto-spectrum of both signals. Spectral coherence relates to TRFs since their Fourier transforms are the cross-spectrum divided by input signal.

Various methods exist for estimating spectral coherence, where the Welch method is the most commonly applied. In this approach, uncorrelated Fourier-based spectrum estimates from multiple data segments (that may overlap) are averaged [17]. Another approach is Thomson’s multitaper method, which multiplies data with various tapering windows before Fourier analysis and subsequent averaging [18–20]. Multitaper spectral estimation is nowadays a commonly applied tool for EEG [21–24]. Multitapers are also applied in EEG coherence estimation for further detection or classification, as is done in this work [15, 25–28].

The statistics in coherence estimation using Welch method [29, 30] as well as the Thomson’s multitaper method [18, 25, 31, 32] has been thoroughly studied. The coherence of 1 signifies a perfect linear connection between signals, while no coupling yields the ideal coherence is 0. Nevertheless, in practice, the lack of infinite data always introduces a significant bias upwards towards positive values in the no coupling case [29]. In comparison, the variance can be assumed to be small when a reasonable amount of data is available [18, 30]. Given the high noise levels in EEG and the relatively small effect of method choice on variance, we specifically focus on other aspects of the estimation methods.

This paper introduces a method for EEG-speech envelope coherence estimation. The method aims to decrease the bias of estimation, while assessing the effectiveness of coherence in analyzing the relationship between speech envelope and EEG responses. With a certain choice in the cross-spectra estimation, as part of the coherence measure, the bias at low linear coupling can be reduced compared to previous methods. Considering that EEG responses are susceptible to random noise, any detected linear coupling is expected to be weak. Real data is employed to evaluate the coherence methods and showcase their potential in decoding auditory attention. Furthermore, since coherence is applied to signals influenced by 1*/f* spectrally shaped noise prevalent in EEG data [33], this study also investigates biases arising from this phenomenon. Coherence “levels out” the slanted noise in the coherence spectrum, peaks that are widened by the taper kernel will have a bias towards higher frequencies. This paper quantitatively analyzes of this bias and its implications for EEG and hearing application. The study’s hypotheses are as follows.

### Two anticipated biases in Speech-EEG Coherence Estimation

1. Due to high level of noise present in EEG data, we anticipate an overall upward bias across all frequencies from all methods. However, we propose that the new method can mitigate this bias. (2) We expect coherence peaks from all multitaper coherence estimates to shift towards higher frequencies, due to the slanted nature of EEG noise. The extent and nature of this shift is hypothesized to be dependent on factors such as the number of tapers, data length, and the shape and power of the noise present in signals.

### Auditory Attention and Hearing Aid Effects Manifested in Speech-EEG Coherence Changes

1. Differences in speech-EEG coherence between attended and ignored speech, specifically within the delta, theta, and alpha frequency bands associated with auditory processing, are anticipated when applying these methods to a population using hearing aids. For these bands, we expect clearer distinctions in coherence estimates with the improved new method, since this improves bias and variance of coherence estimates. (2) Speech-EEG coherence differences are anticipated to be amplified when activated hearing aid noise reduction feature compared to deactivated.

This paper is structured as follows. Firstly, coherence methods are presented in Section 2. Analysis of the positional bias of peaks in the coherence spectrum due to 1*/f* noise is evaluated analytically as well as in simulations. Section 3 outlines the experimental design, pre-processing of data and statistical methods used in real data analysis. Results on real data are presented in section 4, highlighting the capabilities and limitations of coherence methods in auditory attention decoding and in objective evaluation of hearing aid benefits. Conclusions are found in section 5.

## 2 Coherence Estimation

Spectral coherence *C*_*xy*_(*f*) is the measure of the magnitude of a linear coupling between two signals in a system, ranging 0 *≤ C*_*xy*_(*f*) *≤* 1, here defined by its squared form

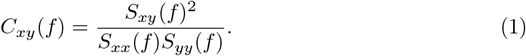

*S*_*xy*_(*f*) is the cross-amplitude spectrum between signals *x*(*n*) and *y*(*n*). *S*_*xx*_(*f*) and *S*_*yy*_(*f*) are the respective auto-spectra. Coherence, as denoted in Eq. (1), is referred to as Magnitude Squared Coherence (MSC).

### 2.1 Magnitude Squared Coherence

Estimating the MSC entails estimating the different spectra according to

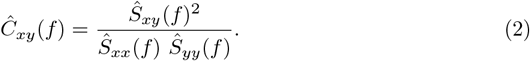

The auto-spectra are estimated through averaging sub-spectra for each pair of L number of data segments and K number of data tapers as

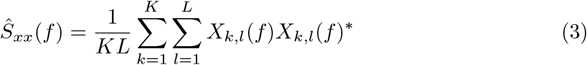

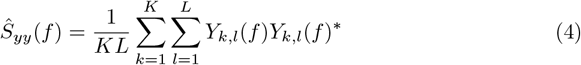

where Fourier transforms of the *l*:th data segments *x*_*l*_(*n*), *y*_*l*_(*n*) and *k*:th tapering window *h*_*k*_(*t*) of respective signal are

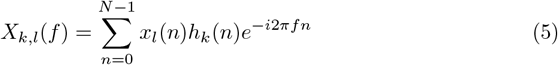

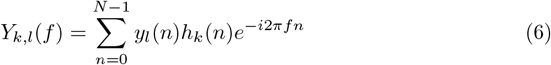

where N is the length of the data segments.

There are multiple options when constructing the cross-spectra from sub-spectra. The different options are all the same in the auto-spectra case, since all subspectra are real and positive. Two options are considered here. The first has been used in [15] in speech envelope to EEG coherence estimation

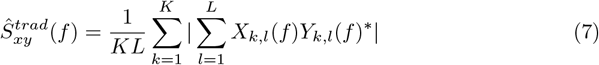

A second option was introduced in our previous work [28] as

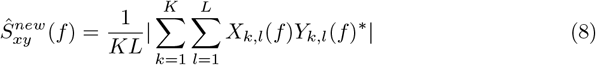

which is shown to decrease the bias upwards in low coherence scenarios. The method in Eq. (8) utilizes more phase information of signals in estimation of the cross-spectrum. A third option where the order of sums in Eq. (7) is reversed, so absolute values of the sum of taper are taken, is not considered here, as *L > K* in most applications.

A common choice, in accordance with the Thomson’s multitaper method, for the set of tapering windows {*h*_*k*_(*t*)} _1,…,*K*_ are the Discrete Prolate Spheroidal Sequences (DPSS), which can be seen in time and frequency domain in Figure 1. These are the windows that maximize the main lobe (or within-band) power relative to broadband power for a normalised frequency bandwidth *W*. They have been shown to have effective variance-reduction characteristics. Although it is argued here that these are suitable choices for tapering windows, the tapering windows can be chosen from another set, with the general methodology staying identical.

**Fig 1.**
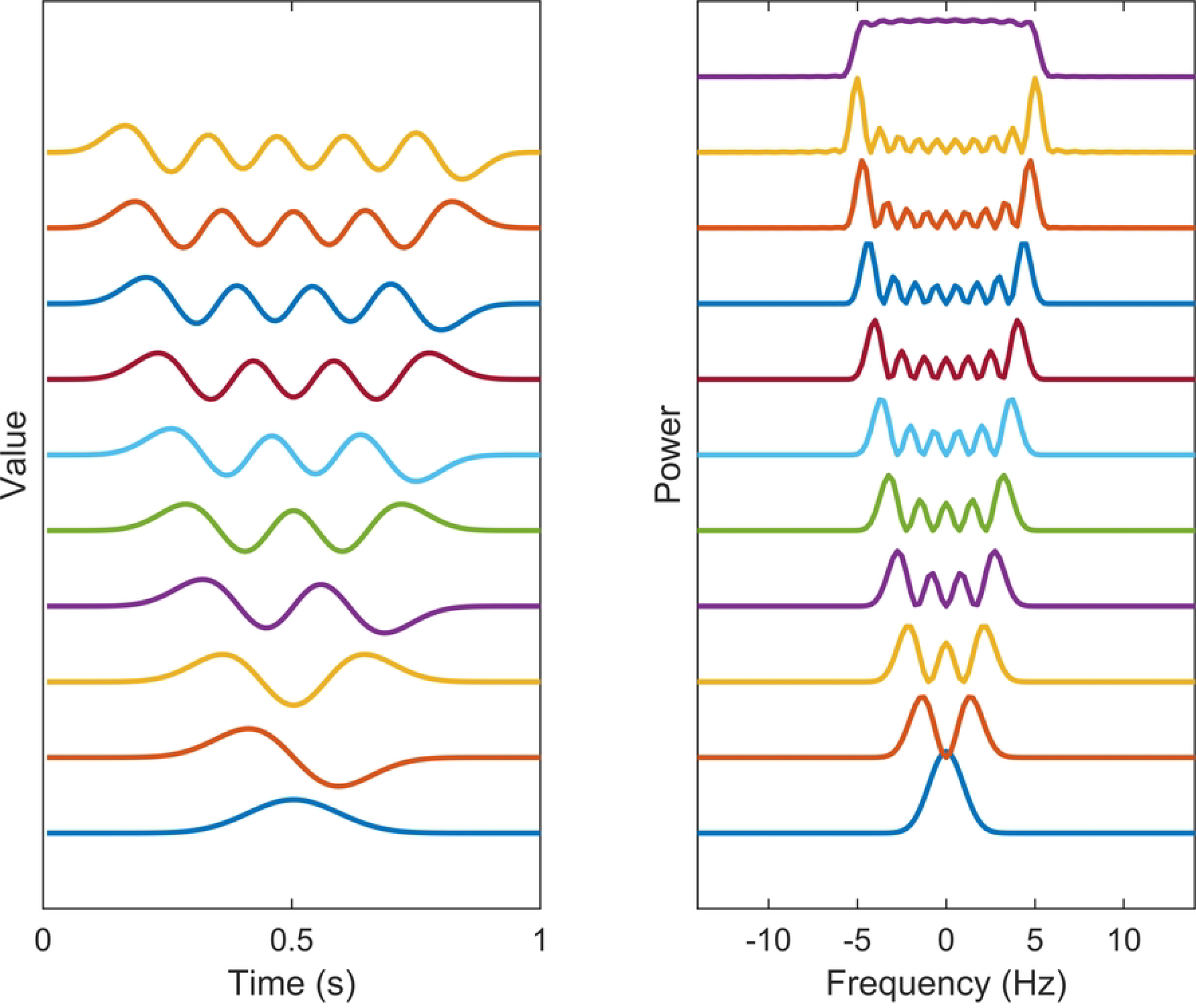
Spectral estimation windows. The temporal representation and power spectra of each of the 10 first DPSS windows used in the multitaper coherence estimation. The sum of the taper kernels is shown above the separate kernels, effectively showing the width of the narrowband leakage effects.

The number of DPSS windows theoretically can be infinite, but using a large amount of tapering windows is not advisable due to its potential to significantly increase the narrowband leakage. This leads to the smearing of the spectrum, making spectral peaks of neighboring frequencies to be indistinguishable from one another. This phenomenon become particularly problematic when estimating coherence in 1*/f* -shaped noise, as discussed in Section 2.2. Instead, a practical approach involves selecting the desired bandwidth of the tapering windows according to the specific application and limit the number of windows accordingly. The DPSS windows are defined in terms of both their shape and the number of windows required to achieve this bandwidth. A common choice for the number of windows is *K* = 2*NW*.

### 2.2 Spectral Positional Bias in EEG-like Noise

In various EEG-related applications, coherence measures are employed to analyze coherence spectra between EEG channels, or between EEG channels and sensory stimuli. This analysis allows the detection of key frequencies showing notable correlation. The identification of these specific correlation frequencies finds applications in tasks like filtering, denoising, and drawing functional insights about the brain.

However, a common challenge faced in visually inspecting the coherence spectra, or any coherence method based on the Fourier transform, is to identify key frequencies in the correlation between channels in the presence of unwanted spectral shifting of relevant coherence energy peaks. This shift is attributed to the presence of slanted 1*/f* noise, indicative of irrelevant brain activity. As the coherence normalizes the base levels of spectrum, the 1*/f* pattern becomes flattened, making it difficult to perceive its impact during peak analysis. This effect is particularly pronounced at lower frequencies, where the 1*/f* noise spectrum exhibits the most prominent degree of gradient.

#### 2.2.1 Approximate Coherence for Sinusoidal Signal

To illustrate the positional bias of peaks in the coherence spectrum when one or more signals are disturbed by 1*/f* noise, we introduce an expression for the coherence of a single frequency coupled model. Although the bias impacts any peaked spectra similarly, a simple case showing the bias involves single frequency oscillations is:

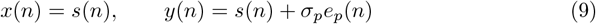

where the signal 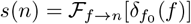 represents a complex sinusoidal and the noise has a spectral distribution *E*[|*ℱ*_*n→f*_ [*e*_*p*_(*n*)]|^2^] = 1*/f*^*α*^, *α >* 0.

In this subsection, it is assumed that 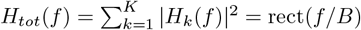, the combined taper kernel, is a box function of energy one and frequency range of *B* = 2*Wf*_*s*_, as shown in [18], where *f*_*s*_ denotes the sampling frequency. The expected value of MSC between *x* and *y* can be approximated by the ratio of smeared cross-spectra and smeared auto-spectra of signals, since these are the expectations of the cross/auto-spectra. This gives a fairly simple expression for the final expectation of the coherence estimate. For *f > B/*2, an initial application of a zero:th order Taylor expansion of the expectation yields

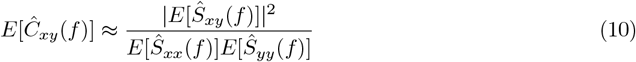

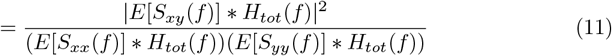

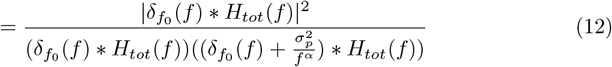

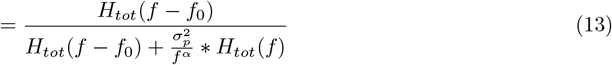

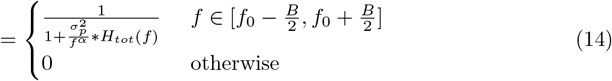

where

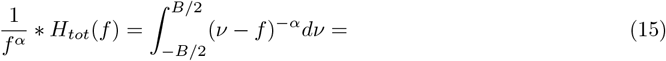

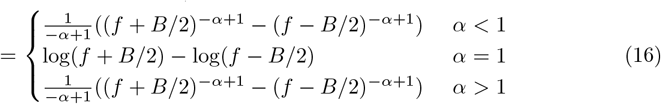

As can be easily observed in Eq. (14), the actual bias beyond the main lobe of the taper kernel is disregarded. In reality, these spectra have some estimated coherence, although lower than the main lobe coherence shown, as discussed in [28]. One can identify this expression as an increasing coherence spectrum within the bandwidth of the tapers used. Perhaps the most frequent choice is *α* = 1, which gives 1*/f* noise. A bias error in Eq. (14) is expected since the approximation is crude in reality. An overall bias upwards is missing, that is observed in the no-coherence case. However, this does not affect the frequency shifting of peaks.

Figure 2 presents a comparison between the empirical expectation of *N*_*real*_ = 10000 simulations of signals according to Eq. (9) to the Eq. (14). Signal parameters were set to *f*_0_ = 10, *N* = 128, sampling frequency *f*_*s*_ = 128, *σ* = [0.5, 1, 5] and *α* = 1, yielding a taper bandwidth *B* = 6 Hz. The simulations demonstrate the bias of the peak clearly in all noise levels of the slanted noise. One can see that the expression follows the expectation of the coherence estimates very well, up to a scale factor.

**Fig 2.**
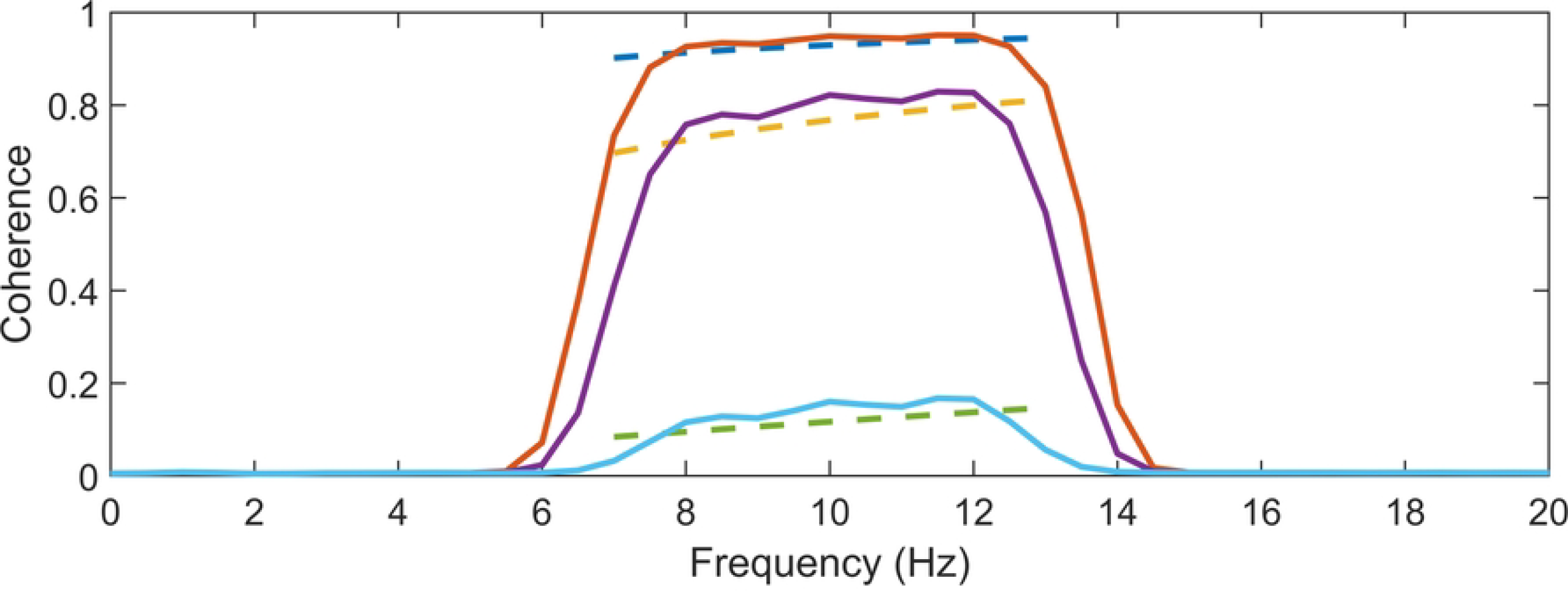
Evaluation of expectation expression. The expression for coherence expectation in dashed lines is shown along the corresponding average coherence of 10000 estimates on a sinusoidal signal in two channels, with added 1*/f* noise in one of the channels. The comparison is made for three different noise levels, *σ* = [0.5, 1, 5], which corresponds the pairs of expectations in ascending order. One can see that the expression estimates the behaviour of the coherence expectation very well, up to a constant. Ideal coherence lies in a single peak at *f*_0_ = 10 Hz.

#### 2.2.2 Effects of Signal and Method Parameters

Characterizing parameter effects on the frequency shifts of peaks in coherence is possible with some a priori knowledge about signals and the coherence estimation method. These shifts depend on the data segment length and the number of tapers used in a multitaper coherence estimation method.

Firstly, the data length *N* modulates the taper kernel’s narrowband width. Decreasing N will linearly increase frequency length of the bias of coherence peaks. Secondly, introducing more tapers in a multitaper scenario will further amplify this bias, although there is a trade-off involving variance reduction with higher *K* values. Simulations are made to visualize this effect. In these simulations, signals are defined as per Eq. (9), with *f*_0_ = 10 Hz. Colored noise *e*_*p*_(*n*) is made by Fourier transforming white noise and then multiplying it by the 1*/f* -spectrum before transforming back. Coherence estimates were computed using the new method on *N*_*real*_ = 10000 generated signals with parameters *N* = 128, *f*_*s*_ = 128, *L* = 50 and varying *K* from 1 to 10. Figures 3a and 3b show the empirical expectation and variance of the coherence estimates across all repetitions. These figures reveal that the coherence peak shifts more in the spectral domain as the number of included window increases. It is worth noting that method parameters, such as data lengths *N* and *K*, are typically chosen in an application and can be adjusted accordingly during analysis.

**Fig 3.**
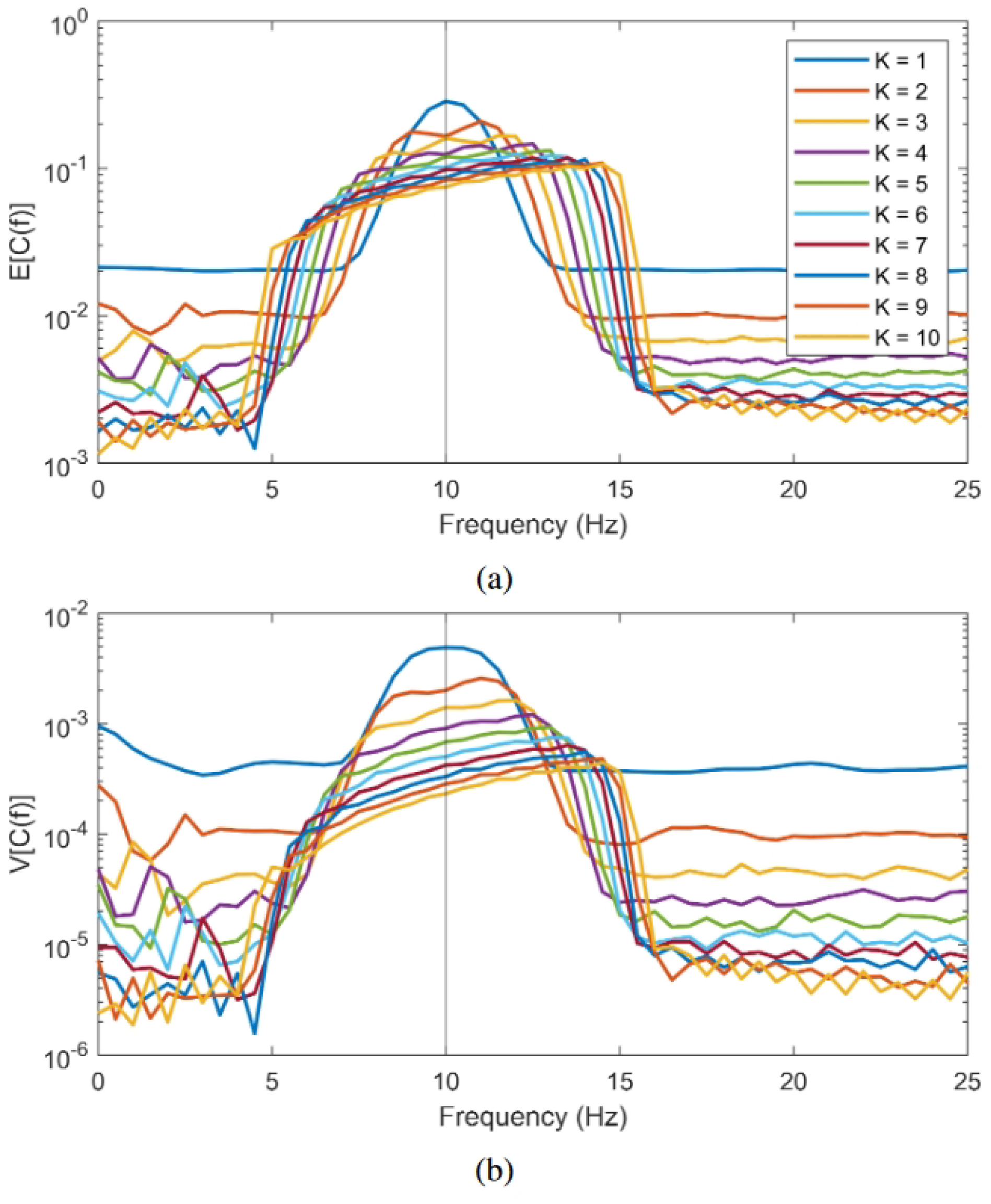
Expectation (a) and variance (b) of coherence estimation using the new method with a varying amount of tapers included. The number of tapers K ranges from 1 to 10. or more blunt peak. This occurs due to the flattening of slope of the 1*/f* noise spectrum at higher frequencies. Additionally, there is often noise in the *x* channel. Parameter effects on the frequency shifting of peaks are consistent when 1*/f* noise (or white) is added to *y*, albeit with an overall lower coherence.

On the other hand, parameters that influence the shifting bias are not as easily quantified prior to analysis. Examples of such parameters include the noise-level of the 1*/f* noise and the spectral position of the coherence peak in question. Referring back to Eq. (14), the SNR of 1*/f* noise doesn’t affect how much the peaks are shifted spectrally; it only impacts the magnitude of the resulting coherence estimate. An exception occurs with very high SNR, where the coherence approaches one, making the slanted peak indistinguishable from a plateau. In this case, the risk of misinterpretation is minimal. Nevertheless, in many applications, coherence values are not substantially large.

It is notable that the true frequency of the peak does not alter how fat peaks are shifted, but that it does influence the slope of the widened band, resulting in a sharper

## 3 Experimental Setup and Statistical Methods

The methods discussed in this study were assessed using an EEG dataset collected from individuals with hearing impairment. This dataset has been previously analyzed in several studies using different analysis approaches [14, 28]. Data collection started 1^st^ of June 2020 and ended on 7^th^ of September 2020. Experimental protocol has been approved by the Science Ethics Committee for the Capital Region of Denmark (journal no. H20028542). The study was conducted according to the Declaration of Helsinki, and all the participants signed a written consent prior to the experiment. More information and details about data can be found in [14].

### 3.1 Participants and Experimental Design

The study included 31 experienced HA users, aged 65.5 years on average, with mild to moderately severe sensorineural hearing loss. All participants were native Danish speakers without a history of neurological disorders, dyslexia or diabetes mellitus. The experiment consisted of 84 trials, with the first 4 trials designated for training and the remaining 80 for testing and analysis. Each trial commenced with a short silence, succeeded by 5 seconds of background noise. Subsequently, 33 seconds of concurrent attended and ignored speech, along with background noise, were presented. Post-trial, participants responded to a two-choice question about the content of the attended speech [12–14].

Speech stimuli were presented through a sound card (RME Hammerfall DSP multiface II, Audio AG, Germany) to 6 loudspeakers (Genelec 8040A; Genelec Oy, Finland). The loudspeakers were positioned at 30^*°*^ (T1-T2), 112.5^*°*^ (B1-B2) and 157.5^*°*^ (B3-B4) azimuth. The audio was played at a 44.1 kHz sampling rate. Attended and ignored speech were presented at 73 dB SPL in T1 or T2, randomized evenly over trials. Background noise was presented at 70 dB SPL in B1-B4, with a mix of 4 talkers in each loudspeaker. T1 and T2 had talkers of opposite sexes presented at each trial. EEG data were recorded at 1024 Hz using a BioSemi ActiveTwo 64-channel EEG system in a 10-20 layout.

Participants wore hearing aids during all trials, split into four sessions of 20 trials each. These sessions were randomly ordered and had different HA noise reduction (NR) settings. Two sessions had NR turned OFF (referred to as HA NR OFF), and two sessions had NR turned ON (referred to as HA NR ON) with two different NR algorithms, as described in [14]. However, these NR algorithms are not discussed in this study.

### 3.2 Preprocessing Steps

The attended and ignored speech signals in each trial were decimated to a sampling rate of 256 Hz, and envelopes were calculated as the absolute value of the analytic version of speech signals, following similar studies [2, 4]. The EEG data were re-referenced to the average of mastoid channels, bandpass filtered between 0.5 and 70 Hz, notch filtered to remove line noise, and resampled to 256 Hz. On average, 0.87 channels per subject were identified as bad channels and were interpolated using neighboring channels.

Independent component analysis [34] was performed to remove components not related to brain activity. On average, 17 components per subject were removed. The analysis involved two speech envelope signals and 64 EEG channels of EEG for each trial.

### 3.3 Statistical Analysis

Statistical p-value testing assessed the significance of method choices (traditional or new coherence estimation method) on differences of speech-EEG coherence between attended and ignored speech. Speech-EEG coherence estimates were computed for each trial using both the traditional method and the new method. This was done using *L* = 33 number of data segments that are *T* = 1 s long, and *K* = 10 data tapers. Coherence values were then averaged across each EEG band (denoted *B*), or retained as estimates for each frequency bin. Bands are defined as delta (1-4 Hz), theta (4-8 Hz), alpha (8-12 Hz), beta (12-30 Hz) and gamma (30-128 Hz). A null hypothesis was formulated, positing that the grand average (over all subjects and trials) of attended speech coherence 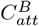 and ignored speech coherence 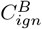 are equal. The alternative hypothesis suggests that 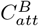 is greater than 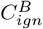 .

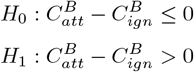

To estimate the null distribution, 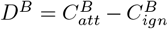, by bootstrap sampling, the sign of *D*^*B*^ was randomized for each subject trial. Next, the mean over all trials and subjects was taken to make one grand average sample from the null hypothesis [17, 23]. This procedure is repeated 500 000 times to approximate the underlying null distribution. Finally, the probability of actual observed coherence grand average difference was derived from the null distribution.

## 4 Results and Discussion

This paper focuses on improving the methods for estimating coherence between speech envelopes and the corresponding EEG responses. This section applies these methods to the experimental data detailed in Sections 3.1 and 3.2. Speech-EEG coherence is shown to be useful in decoding attention in a multi-talker scenario. Furthermore, Section 4.2 provides promising results that highlight speech-EEG coherence as a valuable objective measure for assessing the effects of hearing aid signal processing algorithms on auditory attention.

### 4.1 Separating Attended and Ignored Speech

Figures 4a and 4b show the grand average of coherence estimates using the traditional method and the new method, respectively. Both methods successfully differentiate attended and ignored speech, which is expected due to the success of linear filters in discriminating speakers in multi-talker situations for HI listeners. Notably, the new method provides less noisy coherence estimates across frequencies and exhibits much lower baseline coherence. While both methods show a similar frequency pattern, the traditional method shows two clear coherence peaks, whereas the new method features one more pronounced peak. The new method captures a smaller peak in the 6-9 Hz range for attended speech-EEG coherence, highlighting a difference between speech streams. In contrast, the traditional method lacks this clear pattern.

**Fig 4.**
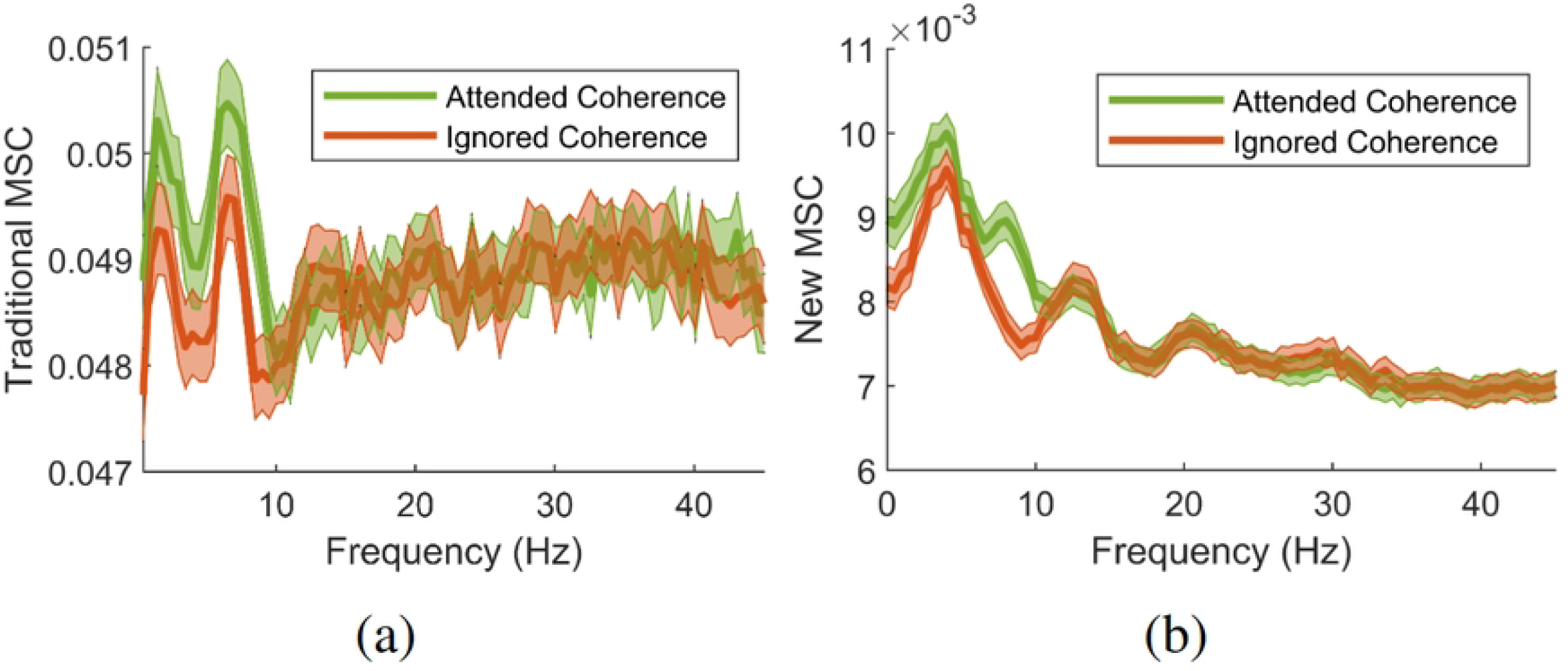
Grand averages of speech-EEG coherence between Attended (green) and Ignored (red) speech envelopes estimated using (a) the traditional method and (b) the new method. Colored areas show the empirical 95% confidence interval of estimates across the spectral range. Note the rather large difference in overall absolute value of estimates between the two methods on the y-axis.

The difference seen in Figure 4 becomes more apparent through grand-level mean testing of the coherence. As detailed in Section 3.3, this approach provides three ways for assessing the significance of speech-EEG coherence differences between attended and ignored speech: at the band level, frequency bin level, and channel level. Figure 5a displays p-values, which evaluate speech-EEG difference between attended and ignored speech, for each frequency bin. Figure 5b shows the significance of coherence differences when coherence estimates are averaged within each EEG band.

**Fig 5.**
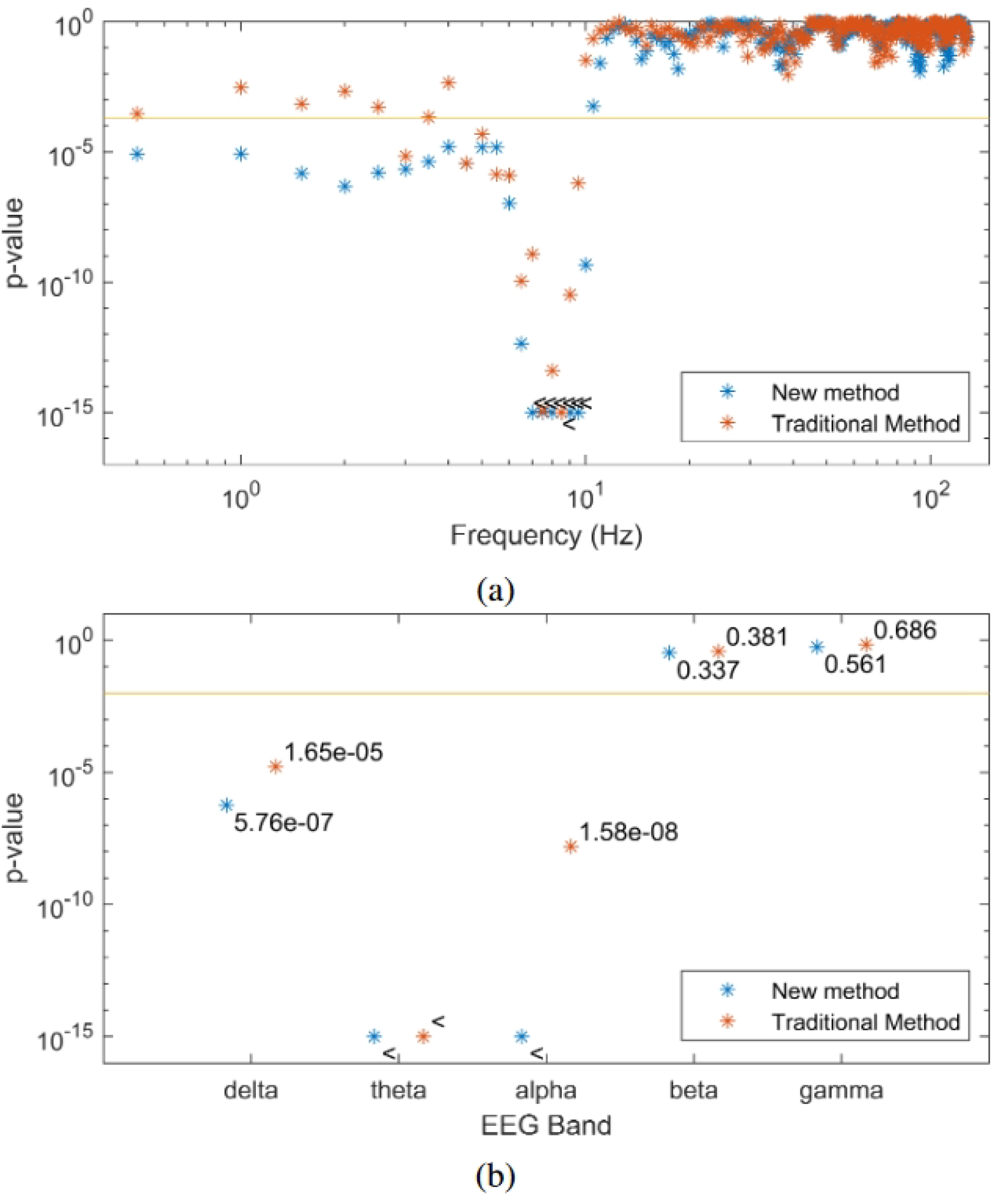
P-values, indicating the difference between observed speech-EEG coherence estimates for attended and ignored speech envelopes, are shown for (a) each frequency bin and (b) each EEG band. Traditional method-derived values are indicated in red, while the values in blue are computed using the new method. The new method consistently yields lower p-values, facilitating stronger statistical testing with reduced data requirements. The 95% significance level of p-values is marked by the horizontal line, with the *<* symbol indicating p-values below the cutoff.

Two main observations can be made. Firstly, significant differences in coherence are observed in the delta, theta and alpha bands. Secondly, within the delta band, multiple individual bins exhibit significance with the new coherence estimation method. This is particularly relevant, since the delta band should indeed exhibit a significant difference between attended and ignored speech envelopes, when estimated using linear filter methods, making the new method superior in this regard to the traditional approach [35, 36]. It is important to note that the new and traditional method yield similar (not significant) p-values for frequencies and bands where a linear relationship between speech envelope and EEG is not anticipated. Also, considering the bias described in Section 2.2, frequency bins showing coherence contain significant information from lower frequencies.

Speech-EEG coherence estimates for the delta, theta and alpha band across each EEG channel were extracted and the grand averages of these were tested, resulting in topographic p-value plots shown in Figure 6. Importantly, enhanced speech-EEG coherence for attended speech envelope compared to ignored speech envelope is more significant when p-values are lower, as depicted by the blue shading in the figure.

**Fig 6.**
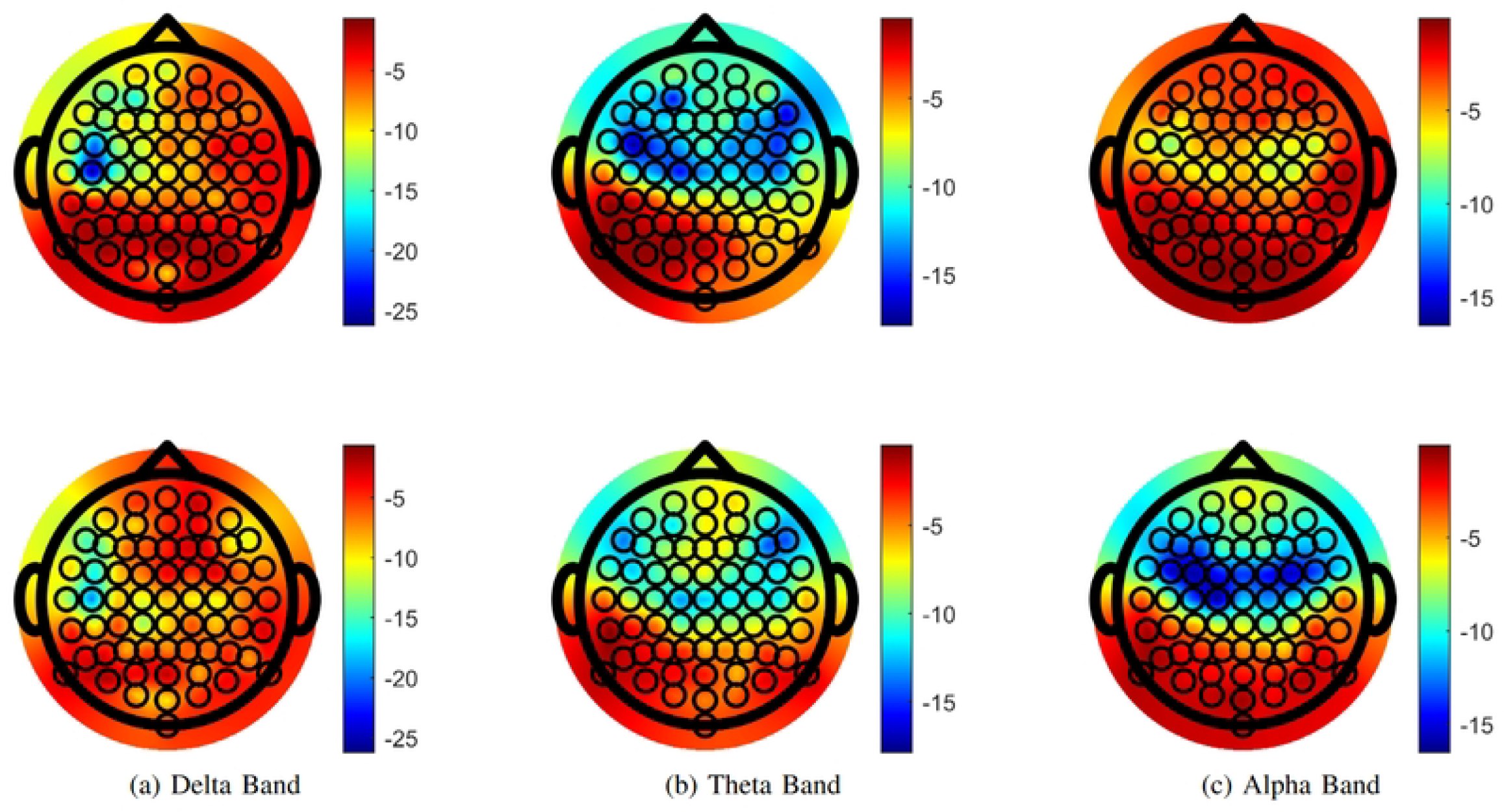
Topographic plots of the log p-values showing significant difference between attended and ignored coherence, (a) the delta band, (b) the theta band and (c) the alpha band. Upper figures show p-values calculated with the traditional method and lower figures are calculated with the new method. Color intensity of p-values is matched over each column for comparison.

Figures 6b and 6c strengthen the evidence for coherence differences in central and frontal regions, consistent with previous research [37–39]. P-values remain low across the parietal channels, in line with prior studies [40]. This pattern is pronounced in the theta and alpha band. Figure 6b and Figure 6c illustrate less noisy patterns over channels when comparing the new coherence estimation method to the traditional one.

### 4.2 Coherence as an Objective Evaluation of HAs

This work suggests using coherence methods to objectively evaluate benefits of signal processing algorithms in HAs in real-life, multi-talker environments. Half of the trials involved participants with the NR algorithm OFF in their hearing aids, while the other half had it ON. After splitting the trials into conditions with HA NR OFF and HA NR ON and conducting identical analysis of attended/ignored speech-EEG coherence differences (now with half the total number of trials for each condition), the resulting p-values are shown in Figure 7a and Figure 7b. P-values are generally lower when HA NR is ON in frequencies showing a linear relationship between speech envelopes and EEG. This suggests a more pronounced distinction between the coherence associated with attended speech compared to coherence associated with ignored speech. It implies that activating HA NR algorithm enhances the separation of attended speech from irrelevant sounds, thus aiding in the auditory attention task.

**Fig 7.**
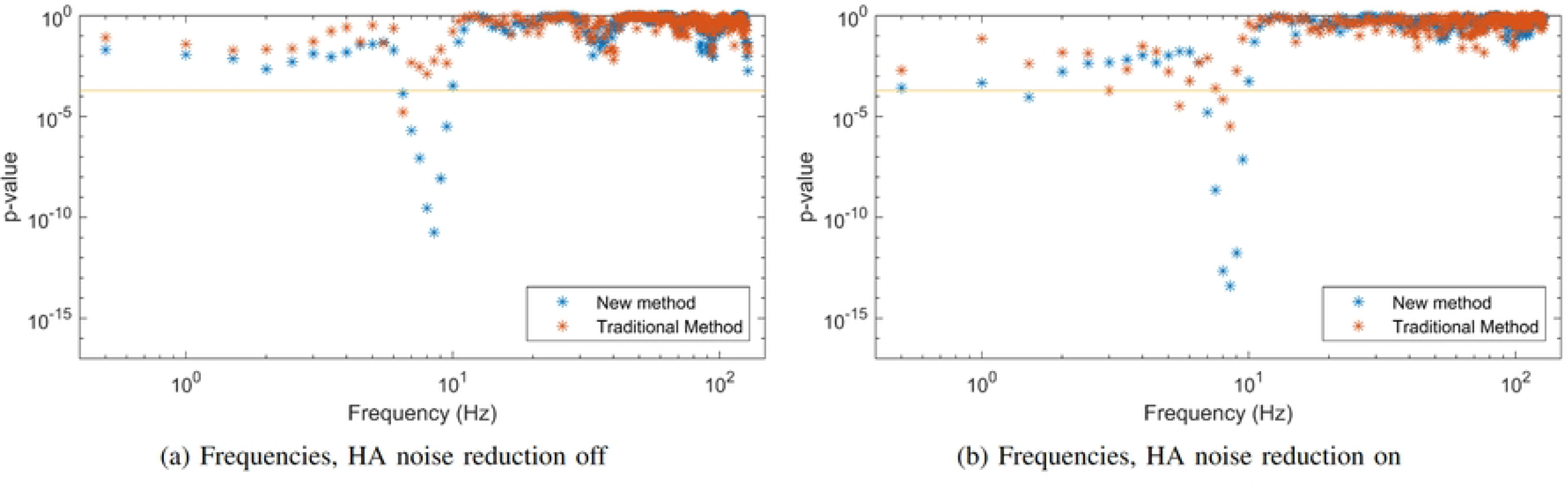
P-values over EEG each frequency bin, for experimental data split between trials with HA NR ON (a) and HA NR OFF (b). Red denotes use of traditional method, and blue denotes instead use of new method. Lower P-values indicate that EEG coherence with attended speech is significantly stronger than with ignored speech. Smaller p-values are generally observed in the HA NR ON condition compared to the HA NR OFF condition, particularly in frequencies of interest.

While beyond the scope of this study, a future research could explore the link between behavioral outcomes and coherence estimates in context of hearing aid signal processing algorithms. Such an investigation could strengthen the proposed objective measure introduced for evaluating the benefits of HA technology. By delving into channels and frequencies of interest, it may demonstrate sensitivity of HAs with less EEG data. This could facilitate clinical adoption of coherence measures as an objective assessment tool for HI and HAs.

## 5 Conclusions

Linear system analysis has previously proven useful for inferring about the connection between speech features, in specific speech envelopes, and measured EEG representing cortical responses. This paper aligns with previous work, and show the functionality of using coherence to decode attention in a cocktail-party environment. Although using higher-order models is useful in their own right, the use of linear models to understand the brain is of importance since their effects on results can be easily quantified.

When interpreting brain electrical activity through spectral aspects of coherence, it is crucial to address two types of biases. Firstly, an upward bias towards higher coherence is analyzed due to responses having a very weak linear connection, showing an improvement with the new method. Moreover, this bias is mitigated by including more trials. Secondly, and of particular significance in EEG applications, there is the shifting of coherence peaks in spectral domain towards higher frequencies. This peak shifting is amplified when widening the effective bandwidth of the spectral estimation technique.

Finally, a novel coherence method was introduced for objectively evaluating the efficacy of hearing aid noise reduction algorithms in realistic listening environments. Observable differences in speech-EEG coherence between attended and ignored speech were observed when the noise reduction feature in hearing aids was activated compared to when it was deactivated. While no rigorous test is presented, the speech-EEG coherence difference between noise reduction schemes remains consistent over frequencies related to the listening task. Further work is needed to explore the suitability of coherence measures for objective evaluation of hearing aids, where parameters such as data amount are taken into account.

